# Acquiring social safety engages oxytocin neurons in the supraoptic nucleus – role of Magel2 deficiency

**DOI:** 10.1101/2024.02.04.578818

**Authors:** Prabahan Chakraborty, Hugo Lamat, Emilie M. André, Pierre Fontanaud, Freddy Jeanneteau

## Abstract

**Introduction:** Exposure to social trauma may alter engagement with both fear-related and unrelated social stimuli long after. Intriguingly, how simultaneous discrimination of social fear and safety is affected in neurodevelopmental conditions like autism remains underexplored. The role of the neuropeptide oxytocin is established in social behaviors, and yet unexplored during such a challenge post-social trauma.

**Methods:** Using *Magel2* knockout mice, an animal model of Prader Willi Syndrome (PWS) and autism spectrum disorders, we tested memory of social fear and safety after a modified social fear conditioning task. Additionally, we tracked the activity of oxytocin neurons in the supraoptic (SON) and paraventricular (PVN) nuclei of the hypothalamus by fibre photometry, as animals were simultaneously presented with a choice between a fear and safe social cue during recall.

**Results:** Male *Magel2* KO mice trained to fear females with electrical footshocks avoided both unfamiliar females and males during recalls, lasting even a week post-conditioning. On the contrary, trained *Magel2* WT avoided only females during recalls, lasting days rather than a week post-conditioning. Inability to overcome social fear and avoidance of social safety in *Magel2* KO mice were associated with reduced engagement of oxytocin neurons in the SON, but not the PVN.

**Conclusion:** In a preclinical model of PWS, we demonstrated region-specific deficit in oxytocin activity associated with behavioral generalization of social fear to social safety. Insights from this study add to our understanding of oxytocin action in the brain at the intersection of social trauma, PWS and related autism spectrum disorders.

## Introduction

Our day-to-day social interactions are complex, requiring discrimination of not only information about the interacting conspecific (age, sex, social status, and so on) but each encounter may also carry a different implicit valence (threat versus mate). Furthermore, several social cues may be present at the same instant (degree of familiarity versus novelty), making such discrimination even more challenging. While neurotypical individuals can appropriately respond to such situations, responses to implicit and explicit social cues are severely compromised in many conditions within the autism spectrum, such as Prader-Willi syndrome (PWS). PWS has been genetically attributed to a lack of expression of 15q11.2-q13 region - containing *MAGEL2* gene - due to paternal deletion or imprinting, or maternal uniparental disomy (1). Apart from well-documented developmental deficits, PWS patients also show a high comorbidity with autism symptoms (2,3) including decreased social responsiveness in non-threatening contexts (4), increased temper outbursts (5–7) and social fear (8). Remarkably, most of these clinical symptoms are well-replicated in *Magel2* knockout (KO) mice, which report impaired response to social novelty following social habituation (9,10) as well as high social fear that is difficult to extinguish (11).

It is interesting to note that *MAGEL2-*deficient patient-derived cell lines express deficient levels of the social neuropeptide oxytocin (OXT) and vasopressin (AVP) (12). Indeed, clinically supplementing OXT by intranasal delivery significantly improves social trust in PWS subjects (13). In *Magel2* KO mice too, OXT treatment early in life prevents social interaction deficits later (9), while AVP can also correct impaired responses to social-novelty in *Magel2* KO adults (10). Remarkably, recruitment of the specific source of OXT and AVP in the brain may be cued to behavioral experience. For instance, OXT neurons in the supraoptic nucleus (SON), but not in the paraventricluar nucleus (PVN), are specifically activated during social fear extinction (11). While we have recently reported that impairment in the SON-lateral septum OXT pathway underlies social fear extinction deficits in *Magel2* KO (11), how the SON neurons *themselves* respond during social fear recall remains unexplored. Intriguingly, common social assays in rodents test response towards a single social cue at one instant. Everyday interactions are complicated by the multiplicity of social cues converging at once, and therefore how *Magel2* KO mice may react to a more ethologically-relevant social challenge remains to be seen.

To address these questions, we designed a novel strategy to condition mice to fear a female, but not a male mouse. We then tested a generalized fear response over days, when unfamiliar males and females were co-presented in a neutral context. Additionally, we tracked the activity of OXT neurons by fiber photometry during these sessions. We believe that observations from this study add new insights into the role of OXT in mediating social behavior within the context of PWS.

## Methods

### Animals

Group-housed animals kept under standard pathogen-free conditions (12/12 light/dark cycle, food and water *ad libitum*) were used in this study. Male *Magel2* KO mice (*Magel2*^tm1.1Mus^/J) alone, or crossed with *Oxt*-Cre (Oxt^tm1.1^(CRE)^Dolsn^/J from Jackson labs and were used in the study. *Magel2* gene is paternally expressed (p) but maternally (m) imprinted, and thus heterozygotes with the paternal null allele (-p) were designated as knockouts (KO). Across experiments, age- and weight-matched heterozygote *Magel2*+m/-p mice were designated as knockouts and *Magel2*+m/+p mice as WT. For stimulus mice, age-matched, non-littermate B6J WT mice were used.

### Stereotaxic injection of viral vectors

6-week old double-transgenic *Magel2* KO or WT mice crossed with Oxt-Cre were unilaterally injected with 500 nL AAV2/5 pGC-CAG-FLEX-jGCaMP7s-WPRE::hGH (CERVO, CA) in either the SON (AP -0.056 cm, ML +/-0.113 cm, DV -0.545 cm) or PVN (AP -0.9cm, ML +/-0.2 cm, DV 0.45 cm). Injections were performed with microinjector-controlled (micro-4, WPI) syringe (nanofil syringe and Nanofil 33G BVLD Needle, WPI). We observed successful recombination leading to expression of GCaMP in 75% OXT neurons (GFP antibody, 1:3000, Abcam, ab13970; NPI-oxytocin antibody, 1:1000, gift from H. Gainer at the National Institute of Health).

### Fibre Photometry

Optic fibre was implanted in either the SON (AP -0.056 cm, ML +/-0.113 cm, DV -0.545 cm) or PVN (AP -0.9cm, ML +/-0.2 cm, DV 0.45 cm). Monofibre optic ferrule (MFC_400/430-0.66_5mm_MF1.25_FLT, DORIC lenses, Quebec, CA) were fixed with dental cement (Paladur, Henry Schein) and animals were given anti-inflammatory medication (meloxicam, 10 mg/kg) and monitored daily in post-surgery isolation for 3 weeks. Calcium (Ca^2+^) signals were explored in freely-moving animals by connecting the implanted fiber optic ferrule to a low-autofluorescence fiber optic patchcord connected to a rotary joint. For photostimulation, connecterized LEDs (CLEDs) emitting sinusoidal signals (GCaMP, 465 nm, and isosbestic, 405 nm) were connected to a fluorescence minicube (consisting of integrated photodetector heads) by a fiber optic attenuating patchcord. Fluorescence acquired through the minicube was digitized and converted by the fiber photometry console to voltage signals, and processed with a low-pass filter (12 Hz). Behavioural data was acquired simultaneously using DORIC Neuroscience studio (Doric Lenses, Quebec, CA). Isosbestic and calcium signals were demodulated in real time with lock-in mode of acquisition. Detection of Ca^2+^ spikes was done using PMATv1-3 setting a threshold of 2.5 SD (14), and custom-written MATLAB code based on (14) were used to calculate positive area under the curve. For this, loss-of-signal artifacts were corrected, followed by smoothing using loess and linearizing using a moving minima algorithm. Next, GCaMP was fit to the isosbestic by using linear regression. The fitted isosbestic was then used to calculate the ΔF/F, by subtracting the fitted isosbestic from the fitted GCaMP, followed by dividing by the fitted isosbestic, and computing a z-score was computed. Behavioral bouts were scored offline during social interactions using BORIS (15).

### Social-fear conditioning

4 weeks after surgery, all animals were handled for a minimum of 3 days, prior to the beginning of the experiment. A separate cohort of animals were also tested without any surgical implant. For baseline test (day -1, **Fig. 1A**), the male test subject was put in the experimental arena (22×30×44cm) to interact during 3 minutes with an unfamiliar male and an unfamiliar female, presented together. Social cues were placed in two diametric ends of the arena in chambers with grills that allowed active physical interaction. 24 hrs later, the test mouse was put in the conditioning arena (11×15×44cm, cleaned with Anios detergent) to freely interact with a novel male stimulus presented alone (M1), during 5 minutes. Subsequently, M1 is replaced with a novel female (F1) on the opposite end of the arena and social fear conditioning was conducted. A scrambled footshock of 0.5mA, 2 sec was given via an electric grid to the test mouse upon each interaction with the female stimulus. The session was terminated after 5 minutes, or if no further social encounter occurred in the 3 minutes following the last shock. During recall on days 1, 2 and 7, the test mouse was introduced into a novel arena (22×30×44cm, patterned walls, different light, cleaned with 10% ethanol) for 5 mins, to allow interaction with a novel male (M2) and a novel female (F2) kept at diametrically opposite corners. All arenas were carefully cleaned between successive animals, and recorded videos were analysed offline with BORIS (15). Heatmaps depicting spatial localization of test subjects in the arena were generated using ezTrack (16).

**Figure 1.**
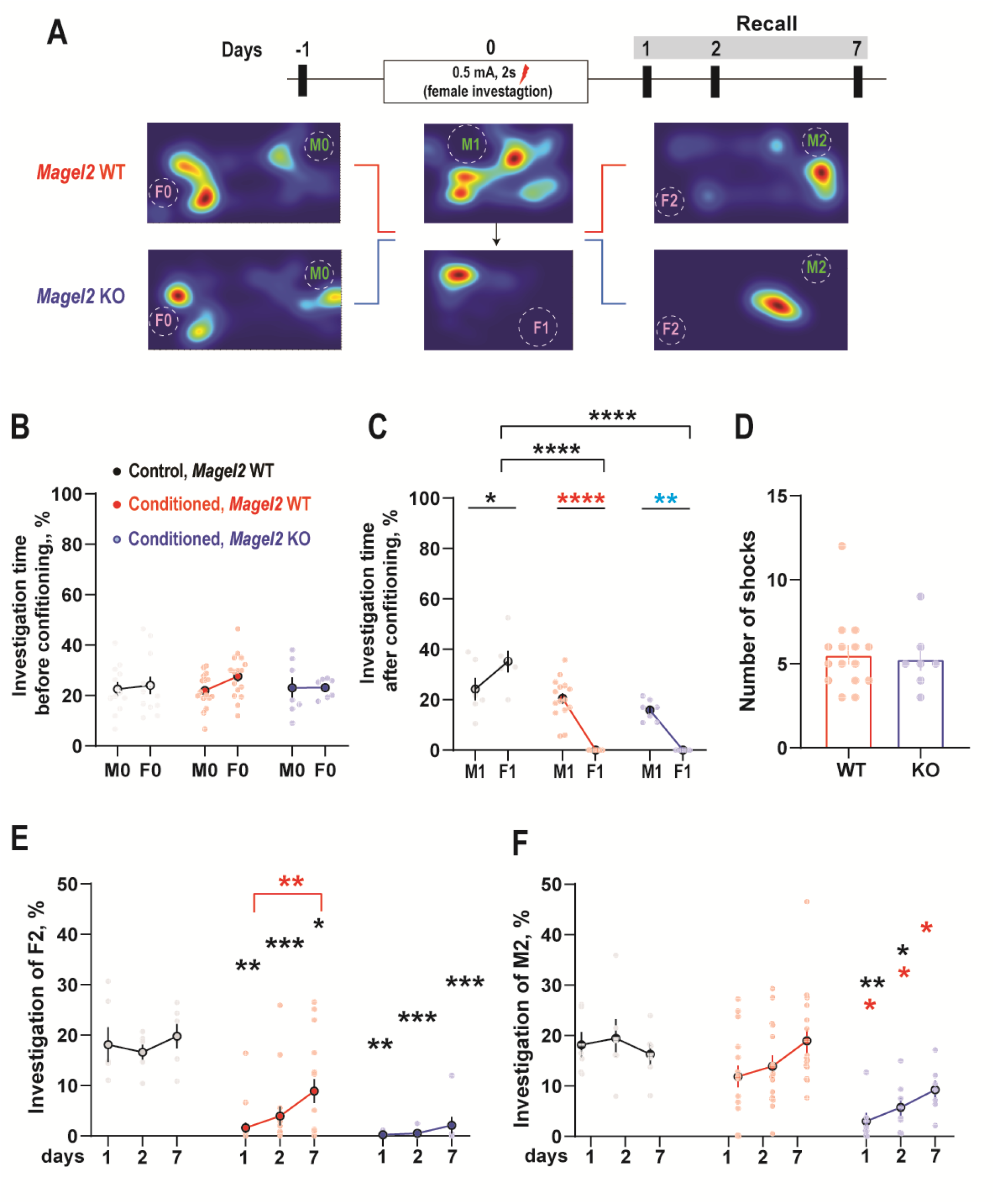
Learning to fear females (social fear cue) generalizes to avoidance of males (social safety cue) in *Magel2* KO mice. (A) Experimental design and timeline. One day before conditioning, test animals interacted with simultaneously presented male and female mice (M0 and F0) in the absence of any shock. During conditioning, *Magel2* KO and WT mice first explored a male stimulus mouse (M1) alone without any shock. This was followed by a session where shock was delivered upon every interaction with a female mouse (F1) (social fear conditioning). 24 h, 48 h and 7 d later, unfamiliar male and female stimulus mice (M2 and F2) were presented at the same time in a novel arena. (B) At baseline before conditioning, all groups interact with M0 and F0 comparably. (Control WT, N = 12, Conditioned *Magel2* WT = 16, Conditioned *Magel2* KO = 7) (C) During social fear conditioning, conditioned *Magel2* WT and KO mice show significantly less interaction with females than males. (Control WT, N = 6, Conditioned *Magel2* WT = 16, Conditioned *Magel2* KO = 7). (E) Following conditioning, *Magel2* KO mice show impaired investigation of F2 over time. Conditioned *Magel2* WT animals show a significant increase at day 7, as compared to day 1. (F) Conditioned *Magel2* KO mice show a robust avoidance of M2 as well, 7 d later. *p<0.05, **p<0.01, ***p<0.001, ****p<0.0001 in post-hoc Sidak’s test. Coloured * indicates comparison to respective group (black, Control, *Magel2* WT, red, Conditioned, *Magel2* WT, blue, Conditioned, *Magel2* KO).

### cFos mapping

On day 7, brain samples were collected 60 minutes after recall via transcardiac perfusion of ice-cold 0.9% NaCl and 4% PFA in deeply anesthetized mice (pentobarbital 50 mg/kg). 40 μm coronal sections was collected with a vibratome, and non-specific sites were blocked prior to immunostaining (non-labeled Fab anti mouse IgG (abliance, 1:500) with 3% normal donkey serum in PBS, 0.1% triton X-100). Primary antibodies (c-Fos: Cell Signaling, mAb 2250, NPI-oxytocin, mouse monoclonal antibodies (H. Gainer, NIH)) were incubated on brain sections for 48 hrs at 4°C, and Alexa-Fluor-conjugated secondary antibodies (1:2,000 ThermoFisher Scientific) were incubated for 2 hrs at 25°C. Sections were mounted with fluoromount and stored at 4°C (ThermoFisher Scientific). Stacks were acquired with an epifluorescence microscope (Imager Z1, Carl Zeiss), and the overlap between cFos and OXT cells counted with FIJI.

### Statistics

Prism 9 (GraphPad, La Jolla, CA) was used for statistical analysis. All data were tested for normality using Shapiro-Wilk and Kolmogorov-Smirnov tests. Photometry data were plotted by events. Normally distributed data were compared with two-way ANOVA, mixed-effects analysis, or unpaired, two-tailed t-test, as applicable. Corresponding post-hoc tests were conducted as necessary. Non-normal datasets were compared using Mann Whitney U-test for two-tailed unpaired comparisons. All data presented are as means ±SEM, N as indicated in figures and statistical tests as indicated in the legends and significance reported as follow: \**p*<0.05, \*\**p*<0.01, \*\*\**p*<0.001, ****p<0.0001. We did not repeat experiments if statistical significance between groups was reached despite small sample size. Details of statistical tests are enlisted in Supplementary Table 1.

## Results

### Social fear conditioning with females generalizes to avoidance of males in *Magel2* KO mice

Studies on social fear in the past have routinely tested fear memory in response to a single social cue. Here, we designed a new task to integrate social fear and safety association based on stimuli-sex **(Fig. 1A)**. In this paradigm, *Magel2* WT and *Magel2* KO mice first investigated an unfamiliar male mouse (M1) in a neutral context **(Fig. 1A**, day 0). Next, they underwent social fear conditioning to an unfamiliar female (F1), such that a scrambled electric footshock was delivered upon each interaction with the female stimuli **(Fig. 1A**, day 0). This created an associative memory that was cued to stimuli-sex, wherein unfamiliar females indicated social fear unlike the unfamiliar male mice that indicated social safety. 24 hours before conditioning, both *Magel2* KO and *Magel2* WT mice explored simultaneously the presented male and female stimuli to a similar extent, indicating no baseline behavioural deficits in absence of shock **(Fig. 1B)**. However, both *Magel2* WT and KO animals investigated the socially fear conditioned female-stimulus significantly less at conditioning **(Fig. 1C)**, during when they also spent more time in the corner away from the social stimulus **(Fig. S1)**. Additionally, an equivalent number of shocks were delivered to both groups **(Fig. 1D)**, suggesting no difference in shock-sensitivity due to genotype.

Next, we tested the memory of social fear/safety over time wherein new sets of unfamiliar male and female mice (M2 and F2) were simultaneously presented in a neutral context on several days after conditioning **(Fig. 1E)**. *Magel2* WT animals explored the female cue significantly less at days 1 and 2, as compared to unshocked controls **(Fig 1E)**. And, 7 days later, however, female-investigation significantly improved from that at day 1 **(Fig. 1E)**. On the other hand, interaction with the male stimulus was not different from unshocked controls, only showing a non-significant trend of increase over time **(Fig. 1F)**. This is in striking contrast to *Magel2* KO animals that robustly avoided the females even at 7 d, suggesting a stronger memory of social fear **(Fig. 1E)**. Even more remarkably, *Magel2* KO avoided interaction with the males as well at all timepoints post-conditioning **(Fig. 1F)**. This indicated that learning to fear females extended to avoidance of males in animals with *Magel2* deficiency.

### Altered calcium dynamics in SON oxytocin neurons of *Magel2* KO during social fear recall

We aimed to track the activity of SON OXT neurons *in* vivo – that have been previously implicated in social fear (11) - during recall of social fear memory over days while in presence of a socially safe cue. Thus, we stereotaxically targeted the SON in oxytocin-Cre mice crossed with *Magel2* KO to inject a Cre-dependent virus expressing GCaMP7s **(Fig. 2A)**, causing successful recombination **(Fig. 2B)**. This allowed us to record calcium-dependent changes in fluorescence from OXT neurons during recall **(Fig. C)**. As compared to conditioned WT animals, SON OXT neurons in *Magel2* KO showed more Ca^2+^ spikes over time, that reached significance 7 d later **(Fig. 2D)**. Interestingly, a cumulative frequency analysis of spike duration revealed greater proportion of shorter-duration spikes in *Magel2* KO at all time-points **(Fig. 2E-2G)**. Yet, Ca^2+^ spike duration significantly decreased in *Magel2* KO animals post-conditioning **(Fig. 2H)**. A decrease in spike duration was coupled with a decrease in inter-spike interval in *Magel2* KO **(Fig. 2I-K)**. While inter-spike interval increased at d 7 in *Magel2* WT, it remained low in mutant animals **(Fig. 2L)**. Together, recall of social fear in *Magel2* KO mice changes the overall Ca^2+^ dynamics of SON OXT neurons.

**Figure 2.**
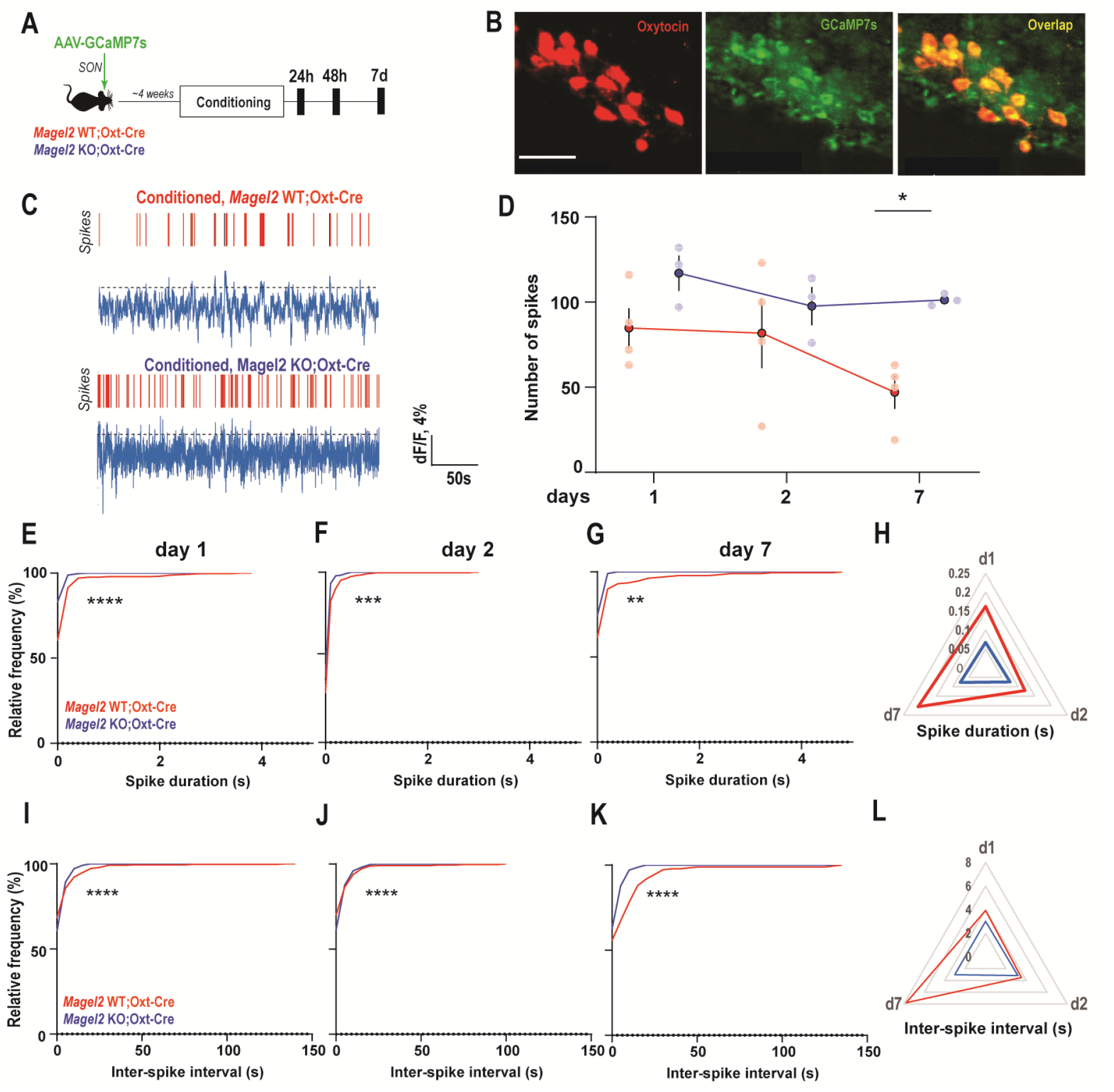
Learning to fear females increases the number, frequency and decreases the duration of Ca^2+^ spikes in SON OXT neurons of *Magel2* KO. (A) Experimental timeline. (B) Representative image showing GCaMP7s expression in SON OXT neurons. Scale bar 50 μm. (C) Representative traces showing Ca^2+^ activity in SON OXT neurons of conditioned *Magel2* WT;*Oxt*-Cre and *Magel2* KO;*Oxt*-Cre mice during recall, 7 d later. Dotted lines indicate threshold for spike determination at 2.5 SD. (D) Social fear conditioning increases the overall number of Ca^2+^ spikes in SON OXT of *Magel2* KO mice. Conditioned *Magel2* WT;*Oxt*- Cre N=4, Conditioned *Magel2* KO;*Oxt*-Cre N=3. * p<0.05, post-hoc Sidak’s test. (E-G) Cumulative frequency of Ca^2+^ spike duration shows a significant left shift post-conditioning. (Conditioned *Magel2* WT;*Oxt*-Cre, d1=338 events, d2=327 events, d7=187 events, 4 animals, Conditioned *Magel2* KO;*Oxt*-Cre, d1=349 events, d2=303 events, d7=292 events, 4 animals) (H) Radar plot showing decrease in spike duration in *Magel2* KO;OXT-Cre mice. (I-K) Cumulative frequency of interval between subsequent Ca^2+^ spikes shows a significant left shift post-conditioning. (Conditioned *Magel2* WT;*Oxt*-Cre, d1, n=334 events, d2, n=323 events, d 7, n=183 events, 4 animals, Conditioned *Magel2* KO;*Oxt*-Cre, d1, n=346 events, d2, n=369 events, d 7, n=300 events, 3 animals). (L) Radar plot showing decrease in inter-spike-interval, indicating increase spike frequency, in *Magel2* KO;*Oxt*-Cre mice. **p<0.01, ***p<0.001, ****p<0.0001 in KS test.

### Reduced engagement of SON OXT neurons in *Magel2* KO *during* investigation of male and female stimuli during recall

The above analysis revealed a generalized change in activity at recall that prompted us to investigate activity of the SON OXT neurons when given the choice to interact with a male and a female. To address this, we looked at changes in fluorescence as the mice actively investigated either male or female stimulus mice before conditioning, and during the days of recall post-conditioning **(Fig. 3A)**. Interestingly, SON OXT neurons had a trend of higher activity (p=0.075) as *Magel2* KO investigated female-stimuli in the absence of any shock **(Fig. 3B1)**. After conditioning, however, a significant reduction in SON OXT activity in *Magel2* KO was observed later at both 1 d **(Fig. 3B2)** as well as 2 d **(Fig. 3B3)** with a slight, qualitative increment observed at 7 d **(Fig. 3B4)**. Overall, we observed a shutdown of SON OXT activity following social fear conditioning that increased with more exploration of females with time in *Magel2* WT, and showed a delayed trend towards increase in *Magel2* KO **(Fig. 3C)**.

**Figure 3.**
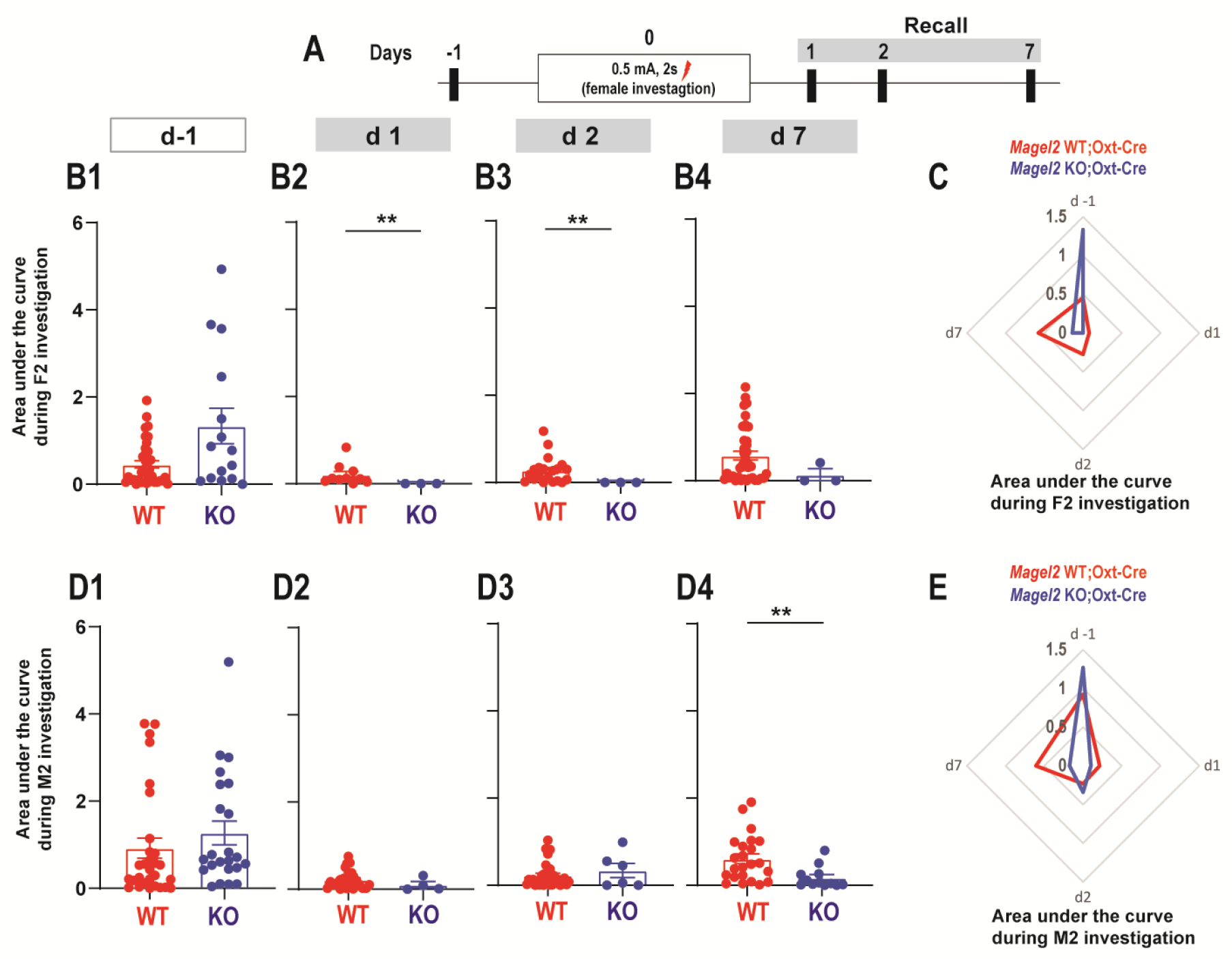
Learning to fear females reduces engagement of SON OXT neurons in *Magel2* KO. (A) Experimental timeline. (B1) *Magel2* KO mice show a non-significant increase in activity during increase of novel female before conditioning. (B2) Post-conditioning, a significant decrease in Ca^2+^ activity is seen during social investigation of novel female mice (social fear cue) at day 1 and (B3) day 2 after conditioning. (B4) Engagement of SON OXT neurons are still low at day 7, although not statistically different (Conditioned *Magel2* WT;*Oxt*- Cre, d 0, N=37, d 1, N=10, d 2, N=20, d 7, N=39, 4 animals, Conditioned *Magel2* KO;*Oxt*-Cre, d 0, N=15, d 1, N=3, d 2, N=3, d 7, N=3, 3 animals). (C) Radar plot depicts trajectory of SON OXT engagement during investigation of female stimulus mice, before and after social fear conditioning. (D1-3) During exploration of an unfamiliar male social-stimuli, difference in SON OXT engagement seen at baseline or at days 1 and 2 after conditioning. (D4) Significant decrease in Ca^2+^ engagement is seen during social investigation of novel male mice at day 7 (Conditioned *Magel2* WT;*Oxt*-Cre, d 0, n=29, d 1, n=26, d 2, n=36, d 7, n=23, 4 animals, Conditioned *Magel2* KO;*Oxt*-Cre, d0, n=4, d 1, n=3, d 2, n=6, d 7, n=12, 3 animals). (E) Radar plot depicts trajectory of SON OXT engagement during investigation of male stimulus mice, before and after social fear conditioning. **p<0.01, in Mann-Whitney U test.

During investigation of male-stimuli at baseline, genotypic difference in SON OXT activity was not observed **(Fig. 3D1)**. In both groups, social fear conditioning significantly reduced SON OXT activity 1 d **(Fig. 3D2)** and 2 d later **(Fig. 3D3)**. However, 7 d later, SON OXT activity in WT animals improved, being significantly higher than *Magel2* KO **(Fig. 3D4)**. This suggested a failure to engaged SON OXT neurons in mutant mice a week after conditioning, in contrast to WT animals **(Fig. 3E)**.

### PVN OXT neurons in *Magel2* KO show changes in calcium dynamics but not differential engagement with overcoming social fear

Beyond the SON, OXT in the brain can originate from other structures such as the PVN, where social neuropeptides have been implicated across a wide range of behaviors. We explored their engagement in the present paradigm, by now targeting the PVN with Cre-dependent GCaMP7s in *Magel2* KO;*Oxt*-Cre mice **(Fig. 4A)**, showing efficient recombination in PVN OXT neurons **(Fig. 4B)**. We next analysed the overall Ca^2+^ spikes during recall **(Fig. 4C)**. Interestingly, we observed a significant decrease in the number of Ca^2+^ spikes during recall **(Fig. 4D)** while the duration of such spikes increased in *Magel2* KO mice **(Fig. 4E)**. A cumulative frequency analysis revealed a significant right-shift in *Magel2* KO animals post-conditioning **(Fig. S2A-C)**. *Magel2* KO mice also showed increased ISI during recalls **(Fig. 4F)**, with a significant right-shift of cumulative frequency post-conditioning **(Fig. S3D-F)**. We next addressed whether PVN neurons differently engaged *during* investigation with the social stimuli. Interestingly, there was no difference in PVN OXT activity due to genotype during female-investigation **(Fig. 4G, Fig. S2G-I)**. While Ca^2+^ dynamics in PVN OXT neurons during male investigation was significantly lower in *Magel2* KO 2 d later **(Fig. 4H, Fig. S2K)**, it did not differ at 1 d and 7 d **(Fig. S2J, L)**. Together, it suggests that social fear conditioning changes the overall Ca^2+^dynamics in *Magel2* KO mice, but its engagement is not associated with overcoming of social fear in either genotype.

**Figure 4.**
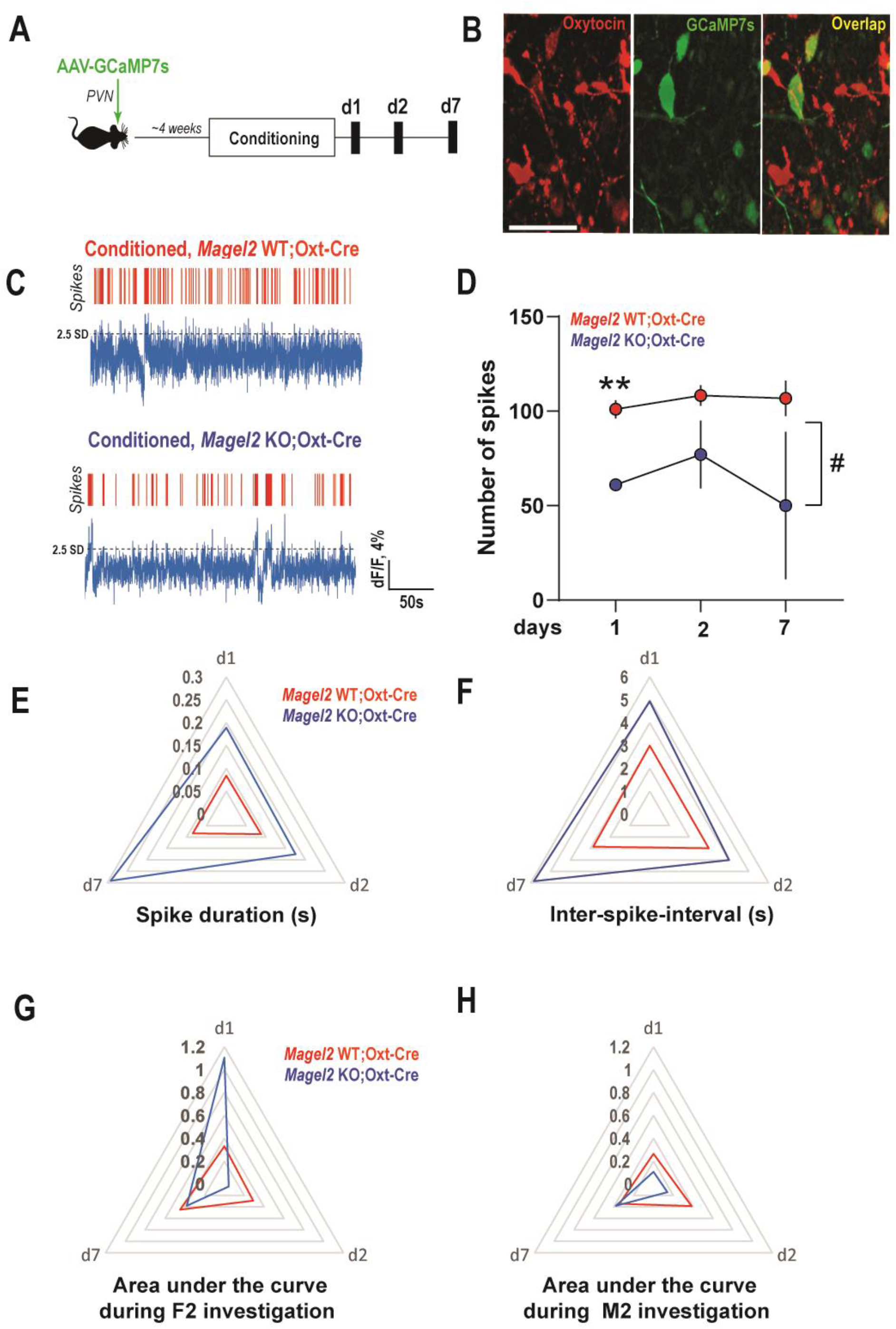
Learning to fear females changes calcium dynamics in PVN OXT neurons in *Magel2* KO, but does not affect their engagement during social investigation. (A) Experimental timeline. (B) Representative image showing GCaMP7s expression in PVN OXT neurons. Scale bar 50 μm. (C) Representative traces showing Ca^2+^ activity in PVN OXT neurons of conditioned *Magel2* WT;*Oxt*-Cre and *Magel2* KO; *Oxt*-Cre mice during recall, 7 days later. Dotted lines indicate threshold for spike determination at +2.5 standard deviation from the mean (SD). (D) Social fear conditioning decreases the overall number of Ca^2+^ spikes in PVN OXT of *Magel2* KO mice. Conditioned *Magel2* WT; *Oxt*-Cre N=5-6, Conditioned *Magel2* KO; *Oxt*-Cre N=2. ** p<0.01, post-hoc Sidak’s test, #p<0.05, main effect of genotype. (E) Post-conditioning, both Ca^2+^ spike duration as well as (F) inter-spike interval increases in *Magel2* KO; *Oxt*-Cre, at all timepoints. (G) Engagement of PVN OXT neurons at days 1, 2 and 7 after conditioning depict during exploration of female stimulus (Conditioned *Magel2* WT; *Oxt*-Cre, d 1, n=10, d 2, n=8, d 7, n=14, 5-6 animals, Conditioned *Magel2* KO; *Oxt*-Cre, d 1, n=2, d 2, n=2, d 7, n=5, 2 animals) (F2) and (H) male stimulus (M2) (Conditioned *Magel2* WT; *Oxt*-Cre, d 1, n=45, d 2, n=110, d 7, n=59, 5-6 animals, Conditioned *Magel2* KO; *Oxt*-Cre, d 1, n=2, d 2, n=8, d 7, n=15, 2 animals).

### Inability to overcome social fear is associated with impaired cFos expression in SON of *Magel2* KO mice

Finally, we mapped the OXT activation patterns of SON and PVN 7 d after conditioning **(Fig. 5B)**, a timepoint where WT animals show a reduction in social fear **(Fig. 1E)**. To that end, we examined the expression of cFos, an immediate early gene produced in response to neuronal activity, in OXT neurons of SON **(Fig. 5B)** and PVN **(Fig. 5C)** 60 minutes after recall. Interestingly, a significant deficit in activation of SON OXT neurons was seen in fear conditioned *Magel2* KO animals as compared to the conditioned WT mice **(Fig. 5D)**. Such a differential engagement was not observed in the PVN **(Fig. 5D)**. In non-shocked controls, OXT neurons in both the WT and KO animals showed no divergent activation as well **(Fig. 5E)**. Thus, activation of OXT neurons that is associated with overcoming social fear in WT is specific to the SON, and such an activation is impaired in the *Magel2* KO model.

**Figure 5.**
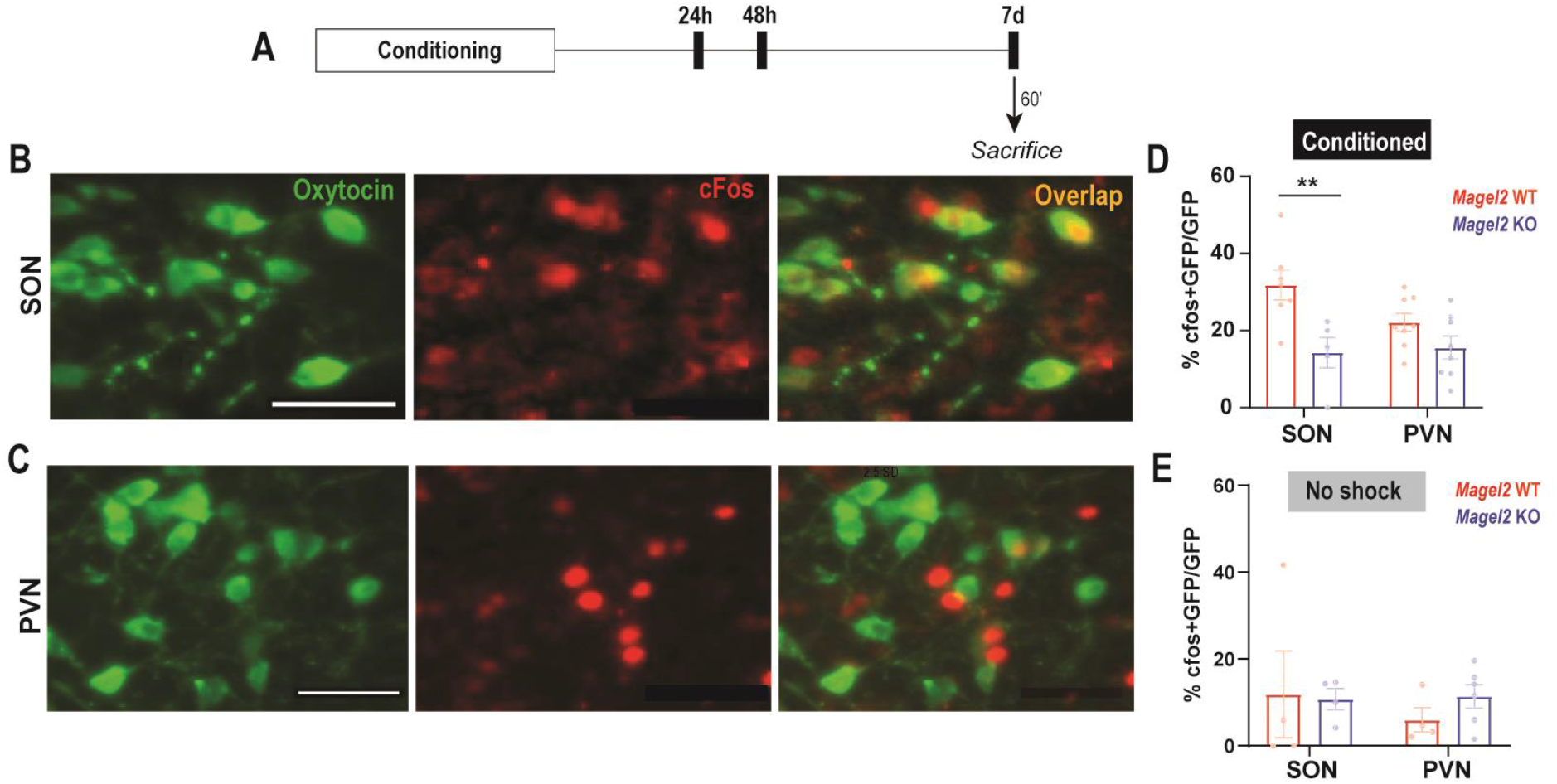
Impaired cFos expression in SON OXT neurons of *Magel2* KO mice, but not PVN, 7 days after conditioning. (A) Experimental timeline. (B) Representative images showing cFos expressing OXT neurons in SON and (C) PVN. Scale bar= 50 μm. (D) *Magel2* KO mice show a significant deficit in cFos expression in SON OXT during recall, 7 d after conditioning, but not in PVN OXT neurons. (*Magel2* WT SON, n=7 roi, N =4 animals, *Magel2* KO SON, n=5 roi, N=3 animals, *Magel2* WT PVN, n=8 roi, N =4 animals, *Magel2* KO PVN, n=8 roi, N=4 animals) (E) No significant change in cFos expression seen in the absence of shock in either group (*Magel2* WT SON, n=5 roi, N =3 animals, *Magel2* KO SON, n=4 roi, N=2 animals, *Magel2* WT PVN, n=4 roi, N =2 animals, *Magel2* KO PVN, n=6 roi, N=3 animals). **p<0.01, post-hoc Tukey’s test.

## Discussion

Using a mouse model of PWS, the present study examined the activity of OXT neurons as conditioned mice recalled social fear while in the presence of a safe social stimulus. Compared to WT, *Magel2* KO mice had similar response during conditioning and equivalent recall of social fear memory 24 h later, in agreement with our recent observations (11). We add that this memory of social fear is persistent and lasts at least up to 7 days after conditioning. However, while WT animals showed a significant increase in exploring the social fear cue at day 7, *Magel2* KO failed to show such an increase. This is reminiscent of the within-session social extinction deficit seen in *Magel2* KO animals (11). We additionally demonstrate that *Magel2* WT animals remember the association of social safety with a male cue and in remarkable contrast, social fear in *Magel2* KO extends to avoidance of socially safe male stimuli. It is interesting to note that impaired discrimination of social salience has been seen with autism – for example, while neurotypical subjects responded faster to a simulated social-hazard, response by autistic subjects was similar towards social and non-social hazards (17). A similar deficit in social salience interpretation may have led to generalization of social fear across stimuli in *Magel2*-deficient animals. It should be noted, however, that mutant mice show a non-significant increase in social safety exploration on day 7. Interestingly in humans, presence of another individual improves the learning of social safety (18), while in rats, learning of safety also helps to overcome threat associated with an avoidance task (19). Together, we predict that a) presence of the social safety cue contributes to the delayed response towards overcoming social fear in *Magel2* KO mice, such that b) overcoming of social fear is preceded by re-learning of social safety in animals with *Magel2* deficiency. One limitation of the present study was that stimuli-females used were not matched to estrus stage. While our results indicate that the behavioural responses are perhaps generalized to female across estrus, it is interesting to note that female mice only in non-receptive phases of the estrus, but not in the receptive phase, show interest towards a same-sex conspecific (20). In male rats, on the other hand, nasal oxytocin abolishes preference towards virgin estrus female rats by diminishing pERK1/2 levels in the SON (21), suggesting a complex relationship between SON OXT action and estrus-dependent social exploration that remains to be explored further. Additionally, previous mechanistic explorations into social fear have used female mice as test subjects (22). Counterbalancing our present results, therefore, it is necessary to investigate social salience in *Magel2*-deficient females trained to avoid males in a future study.

How is neuronal activity of OXT neurons associated with enhanced social fear in *Magel2* KO? To answer this, we recorded activity from OXT neurons in SON and PVN over time in *Magel2* KO and WT animals. In *Magel2*-deficient mice, we observe a shift in Ca^2+^ dynamics leading to an increased number of thinner Ca^2+^ spikes occurring at greater frequency. This is in apparent contrast to previous reports suggesting reduced OXT signaling contributes to heightened social fear (11,22,23). However, given that OXT receptors in *Magel2* KO mice are downregulated (24), it is likely that the increased activity results from homeostatic compensation. A similar compensation is seen in PWS patients – while on one hand they report reduced methylation of OXT-receptor (25) and increased plasma OXT (26), treatment with OXT-receptor agonist improves symptoms of PWS in some clinical studies (27–29) (but not others (30)). Furthermore, spike broadening of OXT neurons as seen here is observed during lactation (31,32), suggesting that reduction of Ca^2+^ spike duration observed in *Magel2* KO mice reflects a physiologically relevant deficit. Indeed, when we specifically examined the activity of SON OXT neurons during social investigation, we observed a reduced engagement of these neurons in *Magel2* KO mice, aligning well with the behavioural deficits observed in these animals **(Fig. 6A)**. Intriguingly, however, SON OXT neurons in *Magel2* KO animals were engaged more during interaction with social cues in the absence of shock **(Fig. 6A)**. Given the positively correlated role of oxytocin in mediating social trust in humans (33), we predict that this aberrant engagement might be necessary to overcome innate threat towards unfamiliar social cues in *Magel2* KO, to enable social approach even at baseline. A subsequent exposure to social fear conditioning shuts down the activity of SON OXT neurons in both genotypes **(Fig. 6A)**, until a divergent trajectory emerges over time during recall. As a result, a progressively more engagement of SON OXT neurons in WT was associated with greater exploration of social fear and socially safe stimuli, in contrast to *Magel2*-deficient animals.

**Figure 6.**
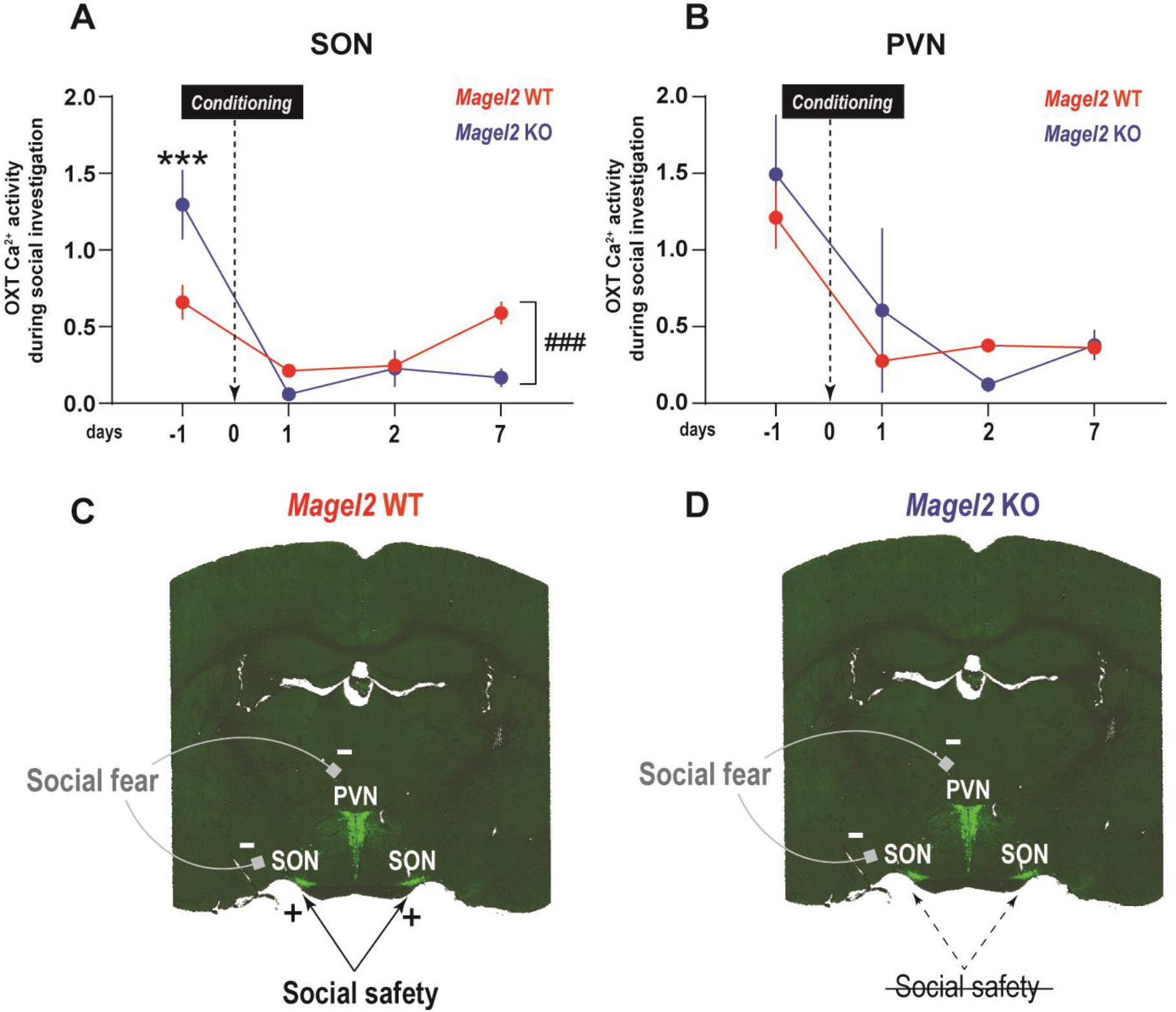
Overcoming social fear and expression of social safety engages SON OXT neurons, and is impaired in *Magel2* deficiency. (A) *Magel2* KO animals show increased engagement of SON OXT neurons during social cue exploration prior to shock-exposure. Social fear conditioning robustly shuts down such engagement in both KO and WT animals. While *Magel2* WT mice can re-engage these neurons over time, *Magel2* KO fails to do so. ***p<0.001, post-hoc Sidak’s test, ###p<0.001, main effect of interaction (genotype x time).(C) No genotypic difference observed in engagement of PVN OXT neurons before or after social fear conditioning. (C) In *Magel2* WT, social fear shuts down OXT neurons in both PVN and SON. Overcoming social fear and expression of social safety selectively reengages SON OXT neurons. (D) In *Magel2* KO mice, social fear also shuts down OXT neuron activity in both SON and PVN, which fail to reengage post-conditioning.

In contrast to SON, cue-specific activity of OXT neurons in the PVN were comparable between WT and *Magel2* KO mice at baseline and although they shut down after conditioning, no significant differences emerged in PVN engagement of OXT neurons over the days of recall **(Fig. 6B)**. This is in agreement with previous reports that show a selective recruitment of SON but not PVN with social fear extinction (11). Interestingly, a reduction in the number and frequency of Ca^2+^ spikes was observed in PVN OXT neurons of *Magel2* KO. This aligns with a previous study reporting decreased firing of PVN OXT neurons in *Magel2*-deficient animals, due to a selective loss of AMPA-current contributing to increased silent synapses (34). We also observed an increase in Ca^2+^ spike duration at the same time point. Interestingly, magnocellular neurons in the PVN, which release OXT (35), have a greater spike half-width than parvocellular neurons (36). Magnocellular OXT neurons projecting to the central amygdala also modulate expression of non-social fear conditioning (37). While we have previously demonstrated that PVN OXT neurons projecting to the lateral septum are not recruited during social fear extinction (11), exploring the role of magnocellular PVN OXT neurons projecting to other targets like the amygdala in social fear needs to be explored in future.

Consistently, mapping of the immediate early gene cFos confirmed the specific engagement of the SON but not PVN OXT neurons in the present study. Increased cFos associated with overcoming of social fear in *Magel2* WT animals 7 days later. It is perhaps intriguing to note that a higher frequency of Ca^2+^ spikes did not lead to more Fos expression in SON OXT neurons of *Magel2* KO. Across brain regions, a complex relationship exists between neuronal activity and cFos expression. In the granule cells, for example, kindled seizures causing action potentials led to cFos expression, but the same stimulation in the substantia nigra failed to do so (38). In the hippocampal CA2, on the other hand, spiking at 0.1 Hz, but not at 50 Hz, was successful in evoking cFos expression, which indicated participation of the neuron in synchronised synaptic activity (39). Interestingly, cFos expression in magnocellular SON OXT neurons is only seen after synaptic activity induced by an OXT-receptor agonist (40). Investigations in the future should focus on understanding the precise role of synaptic inputs in evoking immediate early gene expression in the SON, its lack in the PVN, and their functional interaction with *Magel2* deficiency.

Finally, based on our observations, we propose a model of divergent OXT activation that maps with social fear/safety deficits in *Magel2* KO mice **(Fig. 6C-D)**. In *Magel2* WT animals, exposure to social fear shuts down OXT activity 24 h later in both the SON and PVN **(Fig. 6C)**. However, overcoming social fear to transition towards social safety selectively engages SON OXT neurons **(Fig. 6C)**. What downstream pathways might be activated in such a context? It is tempting to predict that SON OXT neurons might mitigate a hyper-somatostatinergic state in the lateral septum, allowing social investigation even in the face of fear in WT mice (11). In *Magel2* KO, OXT neurons activity shuts down in both the SON and PVN during the initial phase of social fear recall, similar to WT animals **(Fig. 6D)**. However, a failure to engage SON OXT neurons during the behavioural bout accompanies the long-lasting deficit in miscomprehending social safety and thus, the inability to overcome social fear **(Fig. 6D)**. Downstream, a lack of such engagement may fail to inhibit overactivating septal somatostatin neurons, leading to behavioural deficits (11). Additional studies are necessary to identify subsets of SON OXT neurons which specifically encode social safety and fear, and their precise interactions with neuronal ‘functional units’ in the septum that determine exclusive behavioural outputs. In sum, our present demonstration of how OXT neuron activity associates with behavioural approach towards socially salient stimuli provides a fundamental insight into the disease pathology of *Magel2* deficiency. We believe that this set of observations can guide future explorations into treatment strategies in PWS, and other psychiatric disorders with impaired interpretation of social salience.

## Statements

## Acknowledgement

We thank H. Gainer (NIH, USA) for antibodies.

## Statement of Ethics

Experiments were performed according to the Directive by the Council of the European Communities (86/609/EEC) following guidelines from the French Ministry of research and ethics committee for the care and use of laboratory animals (approved protocols APAFIS-5133, 8940, 11468). All efforts were made to minimize animal suffering and to reduce the number of mice utilized in each series of experiments. All experimental procedures in this study were performed between 8:00 and 15:00-hr according to the ARRIVE guidelines.

## Conflict of Interest Statement

The authors declare no conflict of interest.

## Funding Sources

The Fondation pour la Recherche Médicale (FJ, HL) funded this research.

## Author Contributions

Conceptualization: PC, FJ; Methodology: PC, FJ, Investigations: PC, HL, EA; Photometry analysis code: PF, Funding acquisition: FJ; Project administration and supervision: FJ; Writing–original draft: PC, FJ.

## Data Availability Statement

Data will be posted on public repository before publication.

## Notes

### Competing Interest Statement

The authors have declared no competing interest.

## References

1. Dykens EM, Roof E, Hunt-Hawkins H, Dankner N, Lee EB, Shivers CM, et al. Diagnoses and characteristics of autism spectrum disorders in children with Prader-Willi syndrome. J Neurodevelop Disord. 2017 Dec;9(1):18.

2. Fountain M, Schaaf C. Prader-Willi Syndrome and Schaaf-Yang Syndrome: Neurodevelopmental Diseases Intersecting at the MAGEL2 Gene. Diseases. 2016 Jan 13;4(1):2.

3. Kong X, Zhu J, Tian R, Liu S, Sherman HT, Zhang X, et al. Early Screening and Risk Factors of Autism Spectrum Disorder in a Large Cohort of Chinese Patients With Prader-Willi Syndrome. Front Psychiatry. 2020 Nov 26;11:594934.

4. Dimitropoulos A, Ho A, Feldman B. Social Responsiveness and Competence in Prader-Willi Syndrome: Direct Comparison to Autism Spectrum Disorder. J Autism Dev Disord. 2013 Jan;43(1):103–13.

5. Rice LJ, Gray KM, Howlin P, Taffe J, Tonge BJ, Einfeld SL. The developmental trajectory of disruptive behavior in Down syndrome, fragile X syndrome, Prader–Willi syndrome and Williams syndrome. Am J Med Genet. 2015 Jun;169(2):182–7.

6. Singh D, Wakimoto Y, Filangieri C, Pinkhasov A, Angulo M. Guanfacine Extended Release for the Reduction of Aggression, Attention-Deficit/Hyperactivity Disorder Symptoms, and Self-Injurious Behavior in Prader-Willi Syndrome—A Retrospective Cohort Study. Journal of Child and Adolescent Psychopharmacology. 2019 May;29(4):313–7.

7. Tunnicliffe P, Woodcock K, Bull L, Oliver C, Penhallow J. Temper outbursts in Prader-Willi syndrome: causes, behavioural and emotional sequence and responses by carers: Temper outbursts in Prader-Willi syndrome. J Intellect Disabil Res. 2014 Feb;58(2):134–50.

8. Duan Y, Liu L, Zhang X, Jiang X, Xu J, Guan Q. Phenotypic spectrum and mechanism analysis of Schaff Yang syndrome: A case report on new mutation of MAGEL2 gene. Medicine. 2021 Jun 18;100(24):e26309.

9. Meziane H, Schaller F, Bauer S, Villard C, Matarazzo V, Riet F, et al. An Early Postnatal Oxytocin Treatment Prevents Social and Learning Deficits in Adult Mice Deficient for Magel2, a Gene Involved in Prader-Willi Syndrome and Autism. Biological Psychiatry. 2015 Jul;78(2):85– 94.

10. Borie AM, Dromard Y, Guillon G, Olma A, Manning M, Muscatelli F, et al. Correction of vasopressin deficit in the lateral septum ameliorates social deficits of mouse autism model. Journal of Clinical Investigation. 2021 Jan 19;131(2):e144450.

11. Dromard Y, Borie AM, Chakraborty P, Muscatelli F, Guillon G, Desarménien MG, et al. Disengagement of somatostatin neurons from lateral septum circuitry by oxytocin and vasopressin restores social-fear extinction and suppresses aggression outbursts in Prader-Willi syndrome model. Biological Psychiatry [Internet]. 2023; Available from: https://www.sciencedirect.com/science/article/pii/S000632232301661X

12. Chen H, Victor AK, Klein J, Tacer KF, Tai DJC, De Esch C, et al. Loss of MAGEL2 in Prader-Willi syndrome leads to decreased secretory granule and neuropeptide production. JCI Insight. 2020 Sep 3;5(17):e138576.

13. Oztan O, Zyga O, Stafford DEJ, Parker KJ. Linking oxytocin and arginine vasopressin signaling abnormalities to social behavior impairments in Prader-Willi syndrome. Neuroscience & Biobehavioral Reviews. 2022 Nov;142:104870.

14. Bruno CA, O’Brien C, Bryant S, Mejaes JI, Estrin DJ, Pizzano C, et al. pMAT: An open-source software suite for the analysis of fiber photometry data. Pharmacology Biochemistry and Behavior. 2021 Feb;201:173093.

15. Friard O, Gamba M. BORIS : a free, versatile open-source event-logging software for video/audio coding and live observations. Fitzjohn R, editor. Methods Ecol Evol. 2016 Nov;7(11):1325–30.

16. Pennington ZT, Dong Z, Feng Y, Vetere LM, Page-Harley L, Shuman T, et al. ezTrack: An open-source video analysis pipeline for the investigation of animal behavior. Sci Rep. 2019 Dec 27;9(1):19979.

17. Bishop HJ, Biasini FJ, Stavrinos D. Social and Non-social Hazard Response in Drivers with Autism Spectrum Disorder. J Autism Dev Disord. 2017 Apr;47(4):905–17.

18. Pan Y, Olsson A, Golkar A. Social safety learning: Shared safety abolishes the recovery of learned threat. Behaviour Research and Therapy. 2020 Dec;135:103733.

19. Micale V, Stepan J, Jurik A, Pamplona FA, Marsch R, Drago F, et al. Extinction of avoidance behavior by safety learning depends on endocannabinoid signaling in the hippocampus. Journal of Psychiatric Research. 2017 Jul;90:46–59.

20. Chari T, Griswold S, Andrews NA, Fagiolini M. The Stage of the Estrus Cycle Is Critical for Interpretation of Female Mouse Social Interaction Behavior. Front Behav Neurosci. 2020 Jun 30;14:113.

21. Liu XY, Cui D, Li D, Jiao R, Wang X, Jia S, et al. Oxytocin Removes Estrous Female vs. Male Preference of Virgin Male Rats: Mediation of the Supraoptic Nucleus Via Olfactory Bulbs. Front Cell Neurosci. 2017 Oct 23;11:327.

22. Menon R, Grund T, Zoicas I, Althammer F, Fiedler D, Biermeier V, et al. Oxytocin Signaling in the Lateral Septum Prevents Social Fear during Lactation. Current Biology. 2018 Apr;28(7):1066–1078.e6.

23. Zoicas I, Slattery DA, Neumann ID. Brain Oxytocin in Social Fear Conditioning and Its Extinction: Involvement of the Lateral Septum. Neuropsychopharmacol. 2014 Dec;39(13):3027– 35.

24. Gigliucci V, Busnelli M, Santini F, Paolini C, Bertoni A, Schaller F, et al. Oxytocin receptors in the Magel2 mouse model of autism: Specific region, age, sex and oxytocin treatment effects. Front Neurosci. 2023 Mar 14;17:1026939.

25. Heseding HM, Jahn K, Eberlein CK, Wieting J, Maier HB, Proskynitopoulos PJ, et al. Distinct promoter regions of the oxytocin receptor gene are hypomethylated in Prader-Willi syndrome and in Prader-Willi syndrome associated psychosis. Transl Psychiatry. 2022 Jun 10;12(1):246.

26. Johnson L, Manzardo AM, Miller JL, Driscoll DJ, Butler MG. Elevated plasma oxytocin levels in children with Prader–Willi syndrome compared with healthy unrelated siblings. American J of Med Genetics Pt A. 2016 Mar;170(3):594–601.

27. Tauber M, Boulanouar K, Diene G, Çabal-Berthoumieu S, Ehlinger V, Fichaux-Bourin P, et al. The Use of Oxytocin to Improve Feeding and Social Skills in Infants With Prader–Willi Syndrome. Pediatrics. 2017 Feb 1;139(2):e20162976.

28. Tauber M, Mantoulan C, Copet P, Jauregui J, Demeer G, Diene G, et al. Oxytocin may be useful to increase trust in others and decrease disruptive behaviours in patients with Prader-Willi syndrome: a randomised placebo-controlled trial in 24 patients. Orphanet J Rare Dis. 2011;6(1):47.

29. Dykens EM, Miller J, Angulo M, Roof E, Reidy M, Hatoum HT, et al. Intranasal carbetocin reduces hyperphagia in individuals with Prader-Willi syndrome. JCI Insight. 2018 Jun 21;3(12):e98333.

30. Althammer F, Muscatelli F, Grinevich V, Schaaf CP. Oxytocin-based therapies for treatment of Prader-Willi and Schaaf-Yang syndromes: evidence, disappointments, and future research strategies. Transl Psychiatry. 2022 Aug 8;12(1):318.

31. Armstrong WE, Stern JE, Teruyama R. Plasticity in the electrophysiological properties of oxytocin neurons. Microscopy Res & Technique. 2002 Jan 15;56(2):73–80.

32. Stern JE, Armstrong WE. Changes in the Electrical Properties of Supraoptic Nucleus Oxytocin and Vasopressin Neurons during Lactation. J Neurosci. 1996 Aug 15;16(16):4861–71.

33. Tops M, Huffmeijer R, Linting M, Grewen KM, Light KC, Koole SL, et al. The role of oxytocin in familiarization-habituation responses to social novelty. Front Psychol [Internet]. 2013 [cited 2023 Dec 9];4. Available from: http://journal.frontiersin.org/article/10.3389/fpsyg.2013.00761/abstract

34. Ates T, Oncul M, Dilsiz P, Topcu IC, Civas CC, Alp MI, et al. Inactivation of Magel2 suppresses oxytocin neurons through synaptic excitation-inhibition imbalance. Neurobiology of Disease. 2019 Jan;121:58–64.

35. Liu Y, Rao B, Li S, Zheng N, Wang J, Bi L, et al. Distinct Hypothalamic Paraventricular Nucleus Inputs to the Cingulate Cortex and Paraventricular Thalamic Nucleus Modulate Anxiety and Arousal. Front Pharmacol. 2022 Jan 28;13:814623.

36. Chen S, Xu H, Dong S, Xiao L. Morpho-Electric Properties and Diversity of Oxytocin Neurons in Paraventricular Nucleus of Hypothalamus in Female and Male Mice. J Neurosci. 2022 Apr 6;42(14):2885–904.

37. Knobloch HS, Charlet A, Hoffmann LC, Eliava M, Khrulev S, Cetin AH, et al. Evoked Axonal Oxytocin Release in the Central Amygdala Attenuates Fear Response. Neuron. 2012 Feb;73(3):553–66.

38. Labiner D, Butler L, Cao Z, Hosford D, Shin C, McNamara J. Induction of c-fos mRNA by kindled seizures: complex relationship with neuronal burst firing. J Neurosci. 1993 Feb 1;13(2):744–51.

39. Anisimova M, Lamothe-Molina PJ, Franzelin A, Aberra AS, Hoppa MB, Gee CE, et al. Neuronal FOS reports synchronized activity of presynaptic neurons [Internet]. Neuroscience; 2023 Sep [cited 2023 Dec 9]. Available from: http://biorxiv.org/lookup/doi/10.1101/2023.09.04.556168

40. Luckman S, Dyball R, Leng G. Induction of c-fos expression in hypothalamic magnocellular neurons requires synaptic activation and not simply increased spike activity. J Neurosci. 1994 Aug 1;14(8):4825–30.

