## Supplementary figures for "Acquiring social safety engages oxytocin neurons in the supraoptic nucleus – role of Magel2 deficiency"

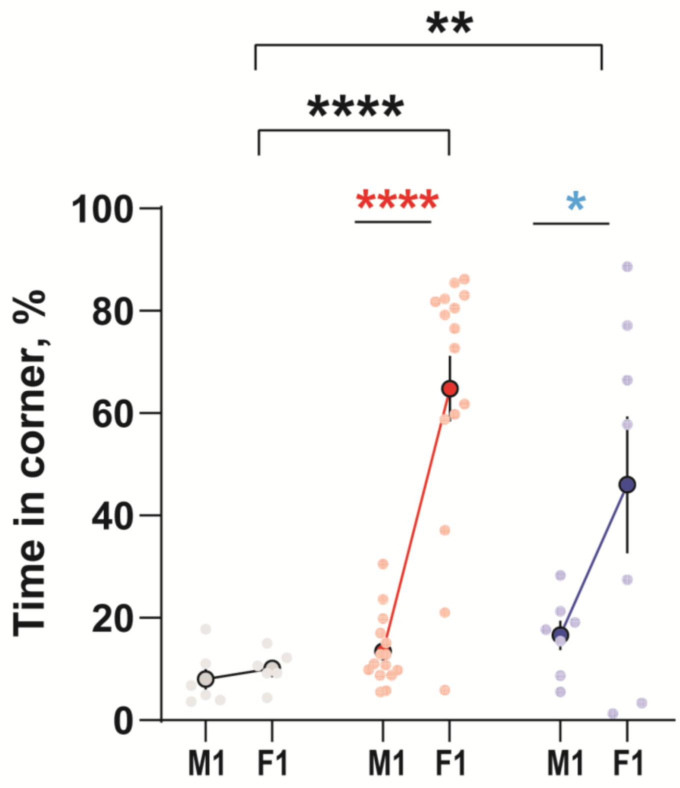


**Supplementary Figure S1 – Time spent in corners during conditioning.** *Magel2* WT and KO mice spend significant time in the corners during social fear conditioning. *p<0.05, **p<0.01, ****p<0.0001 in post-hoc Sidak's test. Coloured * indicates comparison to respective group (black, Control, *Magel2* WT, red, Conditioned, *Magel2* WT, blue, Conditioned, *Magel2* KO).

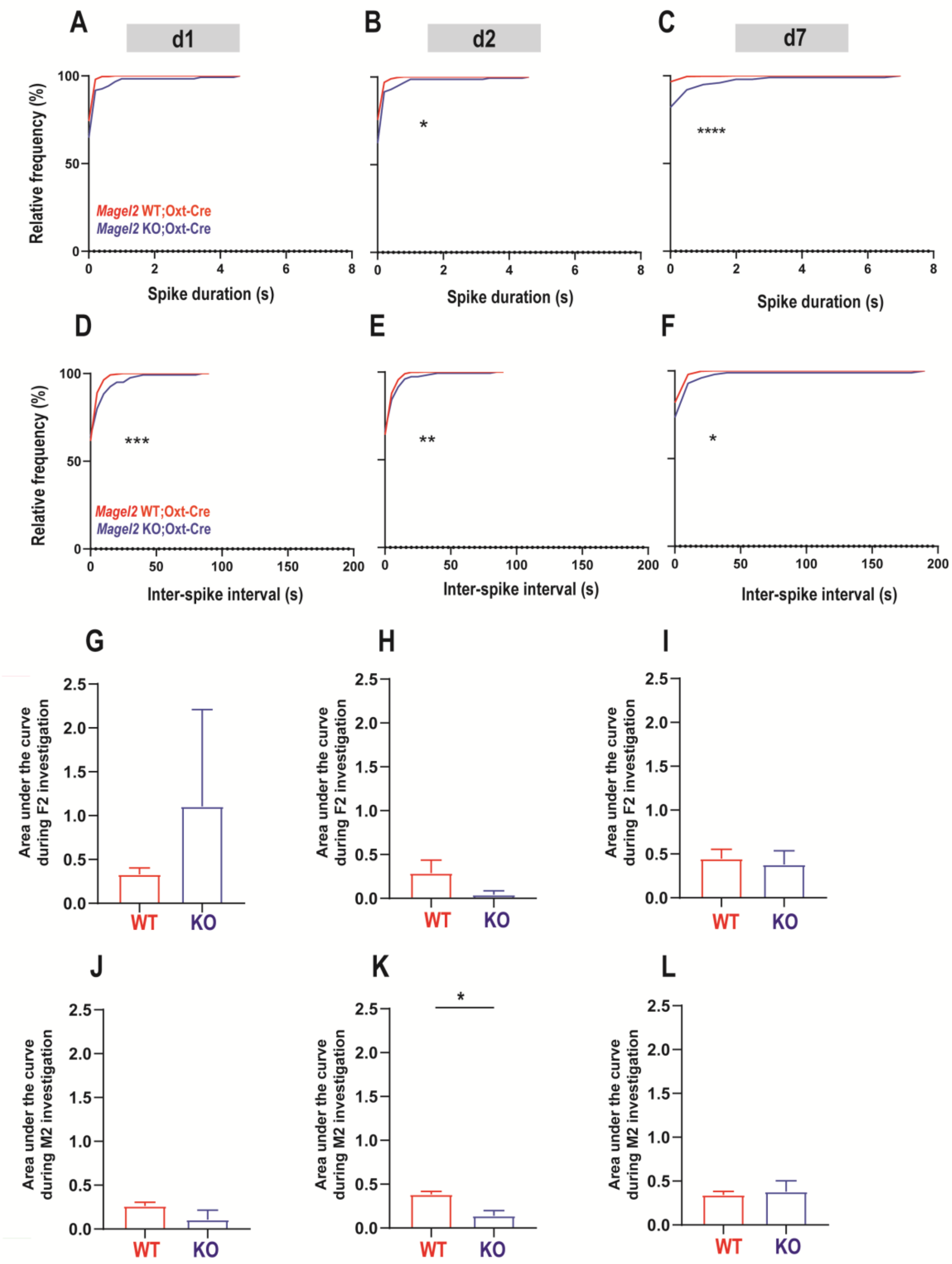


**Supplementary Figure S2 – Altered $Ca^{2+}$ dynamics of PVN OXT neurons following social fear conditioning in *Magel2* KO mice, without significant differences in engagement during behaviour.** (A-C) Cumulative frequency of $Ca^{2+}$ spike duration shows a significant right shift post-conditioning. (D-F) Cumulative frequency of inter-spike interval shows a significant left shift post-conditioning. *p<0.05, **p<0.05, ***p<0.001, ****p<0.0001 in KS test. (G-I) Engagement of PVN OXT neurons are not different during female investigation post-conditioning. (J) Engagement of PVN OXT neurons are not different during male investigation at day 1, (K) transiently increases on day 2, and (L) returns to baseline on day 7. *p<0.05, Mann-Whitney U test.
