## Supplementary Table 1 for "Acquiring social safety engages oxytocin neurons in the supraoptic nucleus – role of Magel2 deficiency"

| Figure reference | Parameter | Statistical test | Statistical details |
|---|---|---|---|
| Fig. 1B | Investigation before conditioning, % | Two-way RM ANOVA | Interaction, F (2, 32) = 0.4441, p=0.65<br>Social stimuli sex, F (1, 32) = 0.8080, p=0.38<br>Genotype, F (2, 32) = 0.4350, p=0.98 |
| Fig. 1C | Investigation during conditioning, % | Two-way RM ANOVA | Interaction, F (2, 25) = 21.01, ****p<0.0001<br>Social stimuli sex, F (1, 25) = 16.71, ***p<0.001<br>Genotype, F (2, 25) = 53.69, ****p<0.0001 |
| Fig. 1D | Number of shocks | Unpaired t-test, two-tailed | t=0.4605, df=15, p=0.65 |
| Fig. 1E | Investigation of F2, % | Mixed-effects analysis | Time, F (1.430, 36.47) = 4.235, *p<0.05<br>Conditioning, F (2, 26) = 21.15, ****p<0.0001<br>Interaction, F (4, 51) = 1.268, p=0.29 |
| Fig. 1F | Investigation of M2, % | Mixed-effects analysis | Time, F (1.859, 47.40) = 1.859, p=0.17<br>Treatment, F (2, 26) = 9.440, ***p<0.001<br>Interaction, F (4, 51) = 1.182, p=0.33 |
| Fig. 2D | Number of spikes | Two-way RM ANOVA | Interaction, F (2, 10) = 1.919, p=0.19<br>Time, F (1.581, 7.907) = 3.716, p=0.08<br>Genotype, F (1, 5) = 5.065, p=0.07 |
| Fig. 2E | Spike duration, d1 | Kolmogorov-Smirnov test | D=.26, ****p<0.0001 |
| Fig. 2F | Spike duration, d2 | Kolmogorov-Smirnov test | D=.16, ***p<0.0001 |
| Fig. 2G | Spike duration, d7 | Kolmogorov-Smirnov test | D=.26, ****p<0.0001 |
| Fig. 2I | Inter-spike interval, d1 | Kolmogorov-Smirnov test | D=.28, ****p<0.0001 |
| Fig. 2J | Inter-spike interval, d2 | Kolmogorov-Smirnov test | D=.21, ****p<0.0001 |
| Fig. 2K | Inter-spike interval, d7 | Kolmogorov-Smirnov test | D=.22, ****p<0.0001 |
| Fig. 3B1 | AUC, female investigation, d-1 | Mann-Whitney U test, two-tailed | Mann-Whitney U=189, p=0.075 |
| Fig. 3B2 | AUC, female investigation, d1 | Mann-Whitney U test, two-tailed | Mann-Whitney U=0, **p<0.01 |
| Fig. 3B3 | AUC, female investigation, d2 | Mann-Whitney U test, two-tailed | Mann-Whitney U=0, **p<0.01 |
| Fig. 3B4 | AUC, female investigation, d7 | Mann-Whitney U test, two-tailed | Mann-Whitney U=23, p=0.09 |

| **Figure reference** | **Parameter** | **Statistical test** | **Statistical details** |
|---|---|---|---|
| Fig. 3D1 | AUC, male investigation, d-1 | Mann-Whitney U test, two-tailed | Mann-Whitney U=244, p=0.1 |
| Fig. 3D2 | AUC, male investigation, d1 | Mann-Whitney U test, two-tailed | Mann-Whitney U=30, p=0.19 |
| Fig. 3D3 | AUC, male investigation, d2 | Mann-Whitney U test, two-tailed | Mann-Whitney U=107, p=0.98 |
| Fig. 3D4 | AUC, male investigation, d7 | Mann-Whitney U test, two-tailed | Mann-Whitney U=56, **p<0.01 |
| Fig. 4D | Number of spikes | Two-way RM ANOVA | Time, F (2, 16) = 0.5707, p=0.36<br>Genotype, F (2, 16) = 0.7960, *p<0.05<br>Interaction, F (1, 16) = 18.58, p=0.52 |
| Fig. 5D | Percentage cFos+ OXT neurons | Two-way ANOVA | Interaction, F (1, 24) = 2.808, p=0.17<br>Brain region, F (1, 24) = 1.634, p=0.21<br>Genotype, F (1, 24) = 13.56, **p<0.01 |
| Fig. 5E | Percentage cFos+ OXT neurons | Two-way RM ANOVA | Interaction, F (1, 14) = 0.4131, p=0.53<br>Brain region, F (1, 14) = 0.2673, p=0.61<br>Genotype, F (1, 14) = 0.1798, p=0.68 |
| Fig. 6A | Area under the curve during social investigation, SON | Two-way ANOVA | Interaction, F (3, 281) = 6.190, ***p<0.001<br>Time, F (3, 281) = 15.06, ****p<0.0001<br>Genotype, F (1, 281) = 0.007346, p=0.93 |
| Fig. 6B | Area under the curve during social investigation, PVN | Two-way ANOVA | Interaction, F (3, 332) = 1.359, p=0.26<br>Time, F (3, 332) = 26.76, ****p<0.0001<br>Genotype, F (1, 332) = 0.5929, p=0.59 |
